# Urea assimilation and oxidation supports the activity of a phylogenetically diverse microbial community in the dark ocean

**DOI:** 10.1101/2024.07.26.605319

**Authors:** Nestor Arandia-Gorostidi, Alexander L. Jaffe, Alma E. Parada, Bennett J. Kapili, Karen L. Casciotti, Rebecca S. R. Salcedo, Chloé M. J. Baumas, Anne E. Dekas

## Abstract

Urea is hypothesized to be an important source of nitrogen and chemical energy to microorganisms in the deep sea; however, direct evidence for urea use below the epipelagic ocean is lacking. Here, we explore urea utilization from 50 to 4000 meters depth in the northeastern Pacific Ocean using metagenomics, nitrification rates, and single-cell stable-isotope-uptake measurements with nanoscale secondary ion mass spectrometry (nanoSIMS). We find that the majority (>60%) of active cells across all samples assimilated urea-derived N, and that cell-specific nitrogen-incorporation rates from urea were higher than that from ammonium. Both urea concentrations and assimilation rates relative to ammonium generally increased below the euphotic zone. We detected ammonia- and urea-based nitrification at all depths at one of two sites analyzed, demonstrating their potential to support chemoautotrophy in the mesopelagic and bathypelagic regions. Using newly generated metagenomes we find that the *ureC gene*, encoding the catalytic subunit of urease, is found within 39% of deep-sea cells in this region, including the Nitrosophaerota (likely for nitrification) as well as thirteen other phyla such as Proteobacteria, Verrucomicrobia, Plantomycetota, Nitrospinota, and Chloroflexota (likely for assimilation). Analysis of public metagenomes revealed *ureC* within 10-46% of deep-sea cells around the world, with higher prevalance below the photic zone, suggesting urea is widely available to the deep-sea microbiome globally. Our results demonstrate that urea is a nitrogen source to abundant and diverse microorganisms in the dark ocean, as well as a significant contributor to deep-sea nitrification and therefore fuel for chemoautotrophy.

## Introduction

Nitrogen (N) is an essential nutrient for all living organisms[1], however, bioaccesible N can be a scarce and therefore limiting element in marine environments[2]. Ammonium and nitrate are among the most important forms of nitrogen in the oceans. While ammonium is assimilable by most microorganisms, nitrate must be enzymatically reduced to ammonium before assimilation, incurring an energetic cost and excluding organisms without this machinery [3, 4]. Ammonium is therefore typically preferred, and is generally scarce (low nM range)[5] while nitrate concentrations can be orders of magnitude higher, especially at depth[6, 7]. Some microorganisms also use inorganic nitrogen as electron acceptors or donors in respiratory processes, increasing the demand for nitrogen in the environment. For example, ammonia can be oxidized to nitrite by chemoautotrophic ammonia-oxidizing archaea (AOA; i.e., Nitrososphaeria, syn., Thaumarchaeota) and ammonia-oxidizing bacteria (AOB)[8, 9]. Nitrogen use in general, and ammonium use in particular, connects closely with carbon cycling, as its availability can influence rates of both heterotrophic[10] and photo/chemo-autotrophic activity [11–13].

Nitrogen dynamics have been studied extensively in the euphotic zone (e.g., [5, 14] ), but less is known about nitrogen cycling in the deep sea, a region increasingly recognized as hosting a diverse, active, and influential microbiome ([7, 15, 16]). Urea, a form of organic nitrogen which can be cleaved enzymatically to create two molecules of ammonia, has been proposed as a key substrate for both anabolism and nitrification in the deep sea [17, 18]. As a source of energy for chemoautotrophy, urea-based nitrification could support organic matter production at depth, thereby ameliorating current discrepancies in the oceanic carbon cycle[19]. However, experimental evidence regarding the abundance [21, 22] and use of urea in the meso- and bathypelagic is still rare or lacking, respectively. Nitrososphaeria-affiliated *ureC* genes and transcripts (encoding urease) have been detected in the epipelagic[17], mesopelagic[19, 23–25] and bathypelagic[26, 27], suggesting the ability of nitrifying archaea to utilize this substrate through the entire water column. Supporting this, urea-based nitrification has been measured at the ocean surface[17], at the base of the epipelagic (at 150 m[28, 29]), and as deep as the mid-mesopelagic (300 m[25])—notably at rates comparable to those for ammonia. Similarly, urea assimilation is extensive in the surface ocean[30], and has been implicated in the mesopelagic based on the observation of urea degradation at rates exceeding calculated N demand for nitrification [31]. However, direct measurements of urea assimilation or oxidation have not been made in the lower mesopelagic or bathypelagic, nor has the prevalence or phylogenetic diversity of organisms containing *ureC* in the aphotic ocean been determined. Therefore, whether the ability to cleave urea is common or rare in the deep sea, taxonomically or numerically, is still unknown, and leaves the accessibility of this potentially large source of nitrogen and energy unconstrained.

In this work, we assessed the role of urea in sustaining microbial biomass production and nitrification from 50 to 4000 m water depth in the northeast Pacific Ocean. We start with an investigation of urea concentrations with depth at six sites across a 300 km transect. At two of these sites, one at the base of the continental slope (“Slope Site”) and one at the far end of the transect (“Open Ocean Site”), we use incubation experiments with ^13^C^15^N-urea and single-cell analysis by nanoscale secondary ion mass spectrometry (nanoSIMS) to determine the proportion of cells assimilating urea-derived nitrogen, and at what rates. We use these same incubations to determine urea- and ammonia-based nitrification rates to assess their role in microbial catabolism throughout the water column. Indeed, although genomic evidence for ammonia-based nitrification at depth is convincing[32, 33], even ammonia-based nitrification has not been experimentally confirmed below the mid-mesopelagic. We furthermore generate thirteen deeply-sequenced metagenomes throughout the Slope and Open Ocean sites, and together with public metagenomes from around the world assess the distribution of the *ureC* gene and the potential role of specific taxa in the utilization and recirculation of urea. Finally, we use the combined ammonium- and urea-based nitrification rates to estimate deep-sea carbon fixation rates, and compare these to estimated rates of sinking particulate organic carbon to estimate the significance of nitrification-based chemoautotrophy at these sites. Together, our lines of inquiry demonstrate the use of urea-derived nitrogen in both microbial anabolism and catabolism in the deep northeastern Pacific Ocean, with implications for nitrogen and carbon cycling globally.

## Material and methods

### Sample collection

Seawater was collected in the northeast Pacific Ocean, off the coast of San Francisco north of Monterey Bay (Figure 1, Fig. S1, Table S1),, onboard the *R/V Oceanus* in March 2017. Seawater was sampled with Niskin bottles at six sites along a 300 km transect (OC1, OC2, OC3, OC4, OC5, and OC6). Samples were collected at 50 m (all sites), 100 m (5 sites), 500 m (5 sites), 1000 m (4 sites), 2000 m (4 sites), 3000 m (4 sites), and 4000 m (1 site), as the water depth allowed. Physicochemical water properties of temperature, conductivity, pressure, and fluorescence were determined with a CTD (SeaBird, USA).

**Figure 1.**
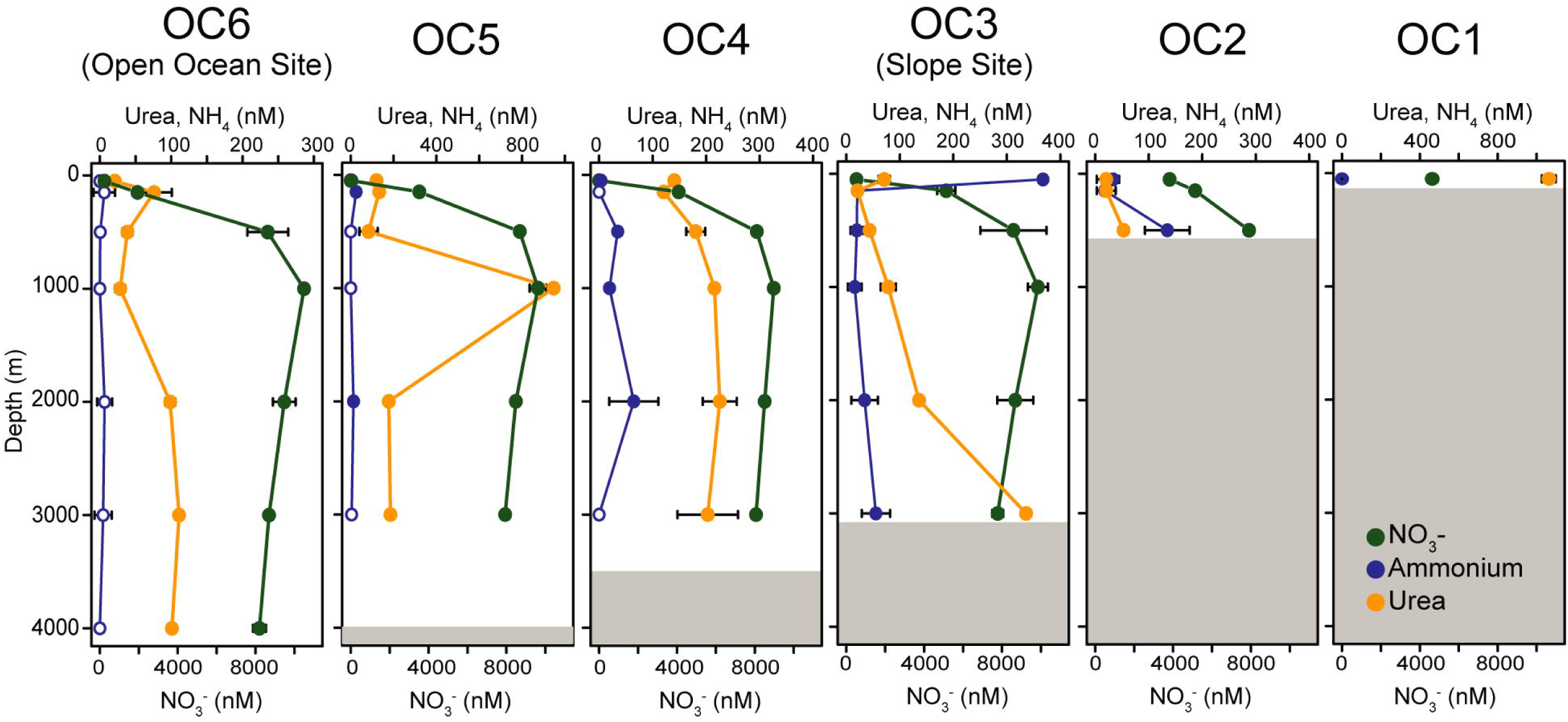
Concentrations of urea, ammonium, and nitrate for each site and depth. Note that the scale for nitrate is different from the other nutrients. Error bars indicate standard deviation of triplicate measurements. Empty dots in ammonium indicate a concentration below the detection limit. Ammonium and nitrate data are re-plotted from Arandia-Gorostidi et al., 2023. Grey area indicates sea-floor depth.

### Quantification of nitrogen species

Water samples for the analysis of nitrogen species were collected from different Niskin bottles in duplicate. Samples were filtered through 0.2μm polycarbonate filters (Isopore) and stored at -80 °C until analysis on shore. Urea concentration was determined following the colorimetric method in Revilla et al. (2005)[34], using 12-hour incubation times at room temperature using replicates. The detection limit was calculated to be 50 nM. Ammonium, nitrate, and nitrite concentrations were previously reported in Arandia-Gorostidi et at. (2023)[7].

### Seawater incubations with stable isotopes

Seawater samples for incubations with stable isotope-labelled substrates were collected at the sampling sites with a maximum depth of 3000 m (OC3, the ‘Slope Site’) and 4500 m (OC6, the ‘Open Ocean Site’). Two sets of incubations were carried out; the first one, previously described in Arandia-Gorostidi et al. (2023) to describe overall microbial activity[7], used 50 nM of ^15^N-labeled ammonium chloride (99% ^15^N, Cambridge Isotope Laboratories, USA), and a second set, newly reported in this study, used 50 nM of ^13^C^15^N-labeled urea (99% ^13^C and 98% ^15^N, Cambridge Isotope Laboratories, USA). Both sets of incubations were conducted as described in Arandia-Gorostidi et al. (2023)[7]. Briefly, seawater samples were incubated in the dark in polycarbonate bottles at 10.5°C (for water samples between 50 m and 150 m depth) or 4 °C (water samples from 500 m to 4000 m). Subsamples from each incubation were fixed using 3% formaldehyde at 0 and 72 hours, and filtered onto polycarbonate filters (25 mm diameter, 0.2 µm pore size; GTTP type, Millipore). Filtered, fixed cells were washed with PBS, 1:1 PBS:EtOH and EtOH before storage at -80 °C for nanoSIMS analysis. Additionally, a portion (15 mL) of the incubated seawater was filtered through polycarbonate filters (25 mm diameter, 0.2 µm pore size; GTTP type, Millipore) into 50 mL Falcon tubes at 0, 24 and 72 hours. The filtrate was stored at -80 °C until nitrification analysis.

### Single-cell isotope uptake by nanoSIMS

Single-cell uptake rates for ^13^C and ^15^N were analyzed by nanoscale secondary ion mass spectrometry (nanoSIMS) using a NanoSIMS 50L (CAMECA, Gennevilliers, France) housed in the Stanford Nano Facility. Analysis conditions are described in the Supplemental Material. Between 61 and 128 cells were analyzed per sample. The isotope images were analyzed using LANS software[35], resulting in the quantitative analysis of isotopic ratios of ^13^C^-12^C^-^/^12^C_2_ ^-^ and ^12^C^15^N^-^/^12^C^14^N^-^. To determine cell-specific isotope ratios, the ^12^C^14^N^-^ channel was used to manually draw regions of interest (ROIs) with outlines just inside the cells. Cells were considered isotopically enriched and therefore consumers of a particular substrate if their isotope ratio was greater than 2 standard deviations above the mean isotope ratio of the 0 h cells from each site[7, 36]. The isotope-based growth (Ka), relative to the initial N and C content, and the single-cell assimilation rates in fg cell^-1^ h^-1^ were calculated following the equations in Stryhanyuk et al. (2018)[37]. The integrated rates for the epi-, meso-, and bathypelagic regions were calculated as described in Arandia-Gorostidi et al. (2023)[7]; multiplying cell density at each depth with the cell-specific assimilation rates (in fg cell^-1^ h^-1^) and the total volume of each region. Statistical differences between the assimilation rates of each substrate were calculated using the Wilcoxon test in R (R version 4.1.3).

### Nitrification rates

Nitrification rates were determined from the rate of production of ^15^N-labeled NOx (NO ^-^+ NO ^-^) in incubations with ^15^N-ammonium and ^15^N-urea at both sites between 150 m and 4000 m depths. 50 m samples could not be analyzed due to the low concentration of NOx at this depth. The ^15^N/^14^N ratio of NOx was determined by isotope ratio mass spectrometry using the denitrifier method[38, 39] in the Stanford Stable Isotope Lab and calibrated using parallel analyses of nitrate isotope reference materials USGS32, USGS34, and USGS35[40]. The nitrification rates were determined using a linear fit of ^15^N-NOx over time in each incubation[41].

### DNA extraction and metagenomic sequencing

Samples for metagenomic analysis were collected at all depths at the Slope Site and the Open Ocean Site. Seawater was filtered through 0.2 µm Sterivex filter units (Millipore, Germany) and flash-frozen in liquid N_2_ immediately after collection. DNA was extracted using the AllPrep DNA/RNA kit (Qiagen, Valencia, CA, USA), following the manufacturer’s protocol. Prior to sequencing, the DNA concentration was measured using the Quant-iT PicoGreen dsDNA Reagent (Invitrogen, Carlsbad, CA, USA). Metagenome sequencing was performed using the NovaSeq S4 PE150 platform at UC Davis sequencing facility (California, USA) with a target sequencing depth of ∼45 gigabasepairs per sample.

### Metagenomic analysis and binning

Paired-end reads were pre-processed with bbduk to remove adapters and to trim low-quality sequences. Trimmed reads were assembled individually using MEGAHIT (v1.2.9[42, 43]). To analyze the distribution and diversity of *ureC* genes within the OC1703 assemblies, as well as those from the public GEOTRACES, TARA Oceans, and Malaspina datasets, we first created a gold standard list of functional *ureC* proteins in cultured marine microorganisms capable of urea degradation (see Table S7). Next, we used this list to identify ureC-encoding contigs using a modification of the PPIT[44] R package. The abundance of *ureC* in each sample was calculated by mapping trimmed metagenomic reads against identified *ureC*-containing contigs with bbmap[45] and quantifying these reads (in terms of reads per kilobase million mapped, or RPKM) with the samtools package[46]. The abundance of *recA*, *amoA*, and nitrate-related genes were calculated using the same approach, though with a modified initial annotation step (Supp. Info). To estimate the propotion of cells containing *ureC*, we normalized its relative abundance to that of *recA*, taking into account the average *ureC* gene copy number per genome observed in the set of metagenome-assembled genomes (MAGs) resolved using methodology described below (1.1 ureC genes/MAG). Additionally, we analyzed *ureC* genetic diversity using a BLAST-based approach that assigns a putative taxonomy to contigs based on the consensus of all genes it encodes (Supp. Info.). The relative abundance of each *ureC*-encoding contig for which taxonomy could be assigned was computed by dividing its sequencing coverage by the total coverage of all *ureC*-encoding contigs in that sample (≥50% breadth). We also subjected the newly-generated OC1703 assemblies to metagenomic binning, forming a set of medium to high quality set of metagenome-assembled genomes (MAGs) ( ≥50% completeness and ≤5% redundancy as determined by CheckM) for downstream analysis (Supp. Info.).[47]

### Estimation of gravitational particulate organic carbon flux

The gravitational Particulate Organic Carbon flux (POC_flux_) at 100 m depth was estimated by multiplying the net primary production by the carbon export efficiency (*e-eff*) calculated according to the following Equation 1[48]:

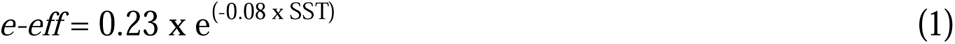

where SST is the Sea Surface Temperature. Satellite-derived net primary production was downloaded from Ocean productivity site (http://sites.science.oregonstate.edu/ocean.productivity/index.php) as 8 days file format treated from the VGPM algorithm[49].

### Statistical Analyses

To test the differences in isotopic assimilation (Figure 2C), the Wilcoxon signed-rank test was used to compare the median assimilation rates between the ^15^N-urea and ^15^N-ammonium incubations at each depth, due to the data distribution not following a normal distribution. Differences were considered significant if p-value<0.05). To test the differences in nitrification rates between the urea and ammonium incubations, we used a t-test to compare the mean rates of each incubation at a given depth. A comparison of the change of ureC relative abundance with depth was performed by ANOVA analysis with the Tukey’s honest significant difference test correction. All the statistical tests were performed in R (R version 4.3.1).

**Figure 2:**
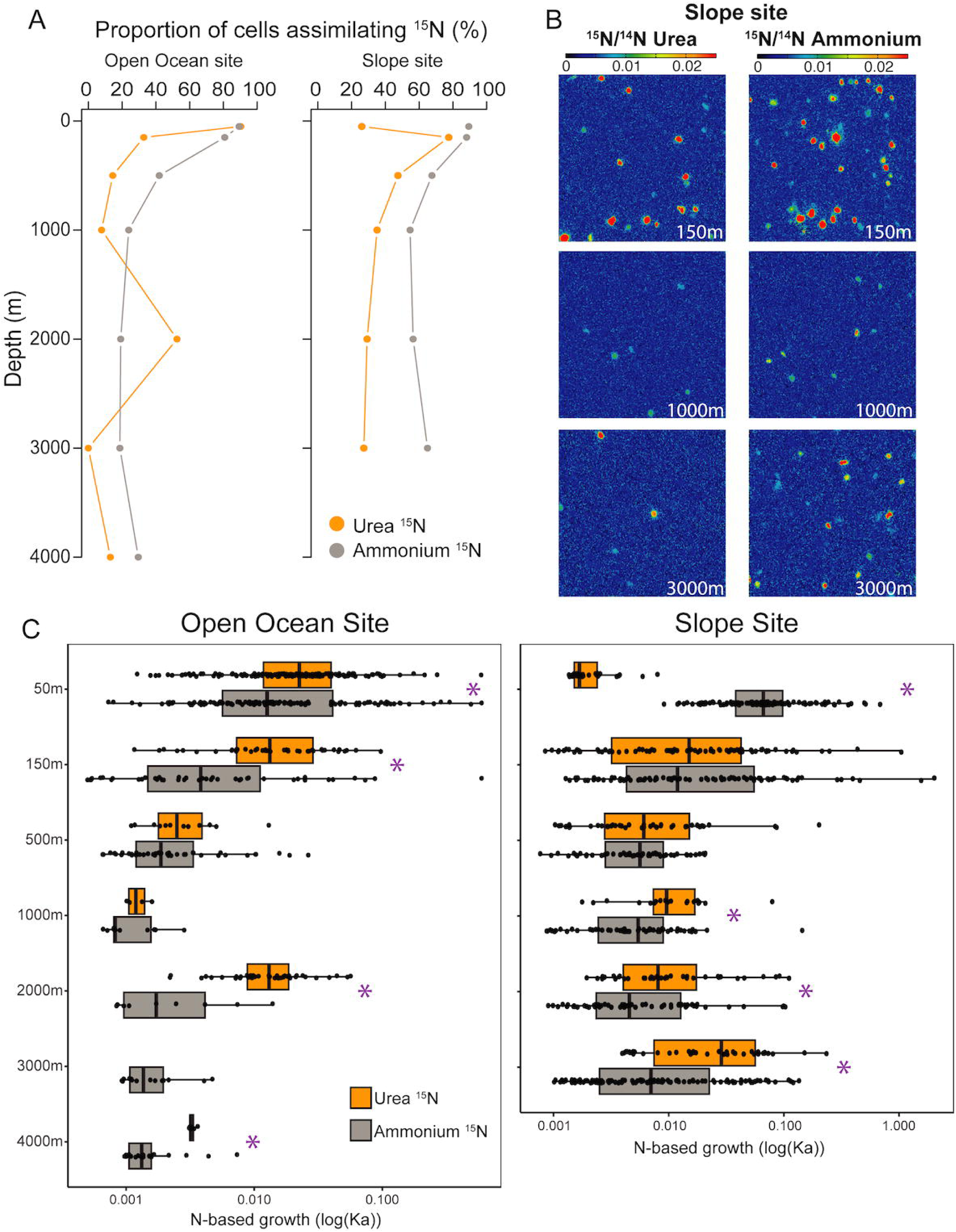
Single-cell assimilation of urea-derived N, compared to ammonium. **(A)** The depth profile plots show the proportion (%) of cells that assimilated nitrogen from urea or ammonium at each depth of the Open Ocean and Slope Sites. **(B)** NanoSIMS images of cells showing assimilation of ^15^N from urea and ammonium in incubations of seawater from the Slope Site. The color scale indicates the ^12^C^15^N^-^/^12^C^14^N^-^ ratio for the analyzed areas. (C) Boxplots represent the N-based growth (Ka) in logarithmic scales for individual cells from both urea- and ammonium-amended incubations at each depth of the Open Ocean Site (left) and Slope Site (right). Purple asterisks indicate that the difference between ammonium and urea rates are significant (ANOVA, p-value<0.05).

## Results

### Quantification of nitrogen species

Urea was detected in all samples, ranging from 21 nM at 50 m water depth at the Open Ocean Site (OC6, 281 km from shore) to 1.1 µM at 50 m water depth at OC1, the most coastal site (14 km from shore) (Figure 1). Urea concentrations were highest below the euphotic zone, and generally higher than those of ammonium. The physicochemical analysis of the sampling sites and the concentration of ammonium and nitrate are described in Arandia-Gorostidi et al. (2023)[7].

### Single-cell urea uptake with depth

Microbial assimilation of urea-derived N was detected in all depths investigated at both the Slope and Open Ocean sites except one (Open Ocean site, 3000 m water depth) (Figure 2). The proportion of cells assimilating urea-derived N as well as the magnitude of incorporation (per cell and the average for all cells assimilating it) was determined and compared to the values previously determined for ammonium assimilation in the same samples[7]. In the Open Ocean site, the highest proportion of cells incorporating urea-derived ^15^N was found at 50 m (90%; Figure 2A), and was almost exactly the same as for ammonium (89%). The proportion of cells incorporating urea-derived N generally decreased with depth, similar to the trend for ammonium, with an exception at 2000 m where the proportion spiked to 53%. The trend in the Slope Site was different from the Open Ocean site, with the lowest proportion of cells incorporating urea-derived N at 50 m (26%) and the highest at 150 m (78%). The proportions in the meso- and the bathypelagic region remained relatively high (37% in average) at the Slope Site, higher than at the same depths at the Open Ocean site (18% on average). The trends in the average N-assimilation rates from urea followed the same pattern as the trends in the portion of cells assimilating urea (Figure 2C). Overall, the Slope Site had a higher proportion of cells assimilating urea-N and at higher rates than at the Open Ocean Site.

On average, urea-N was assimilated at higher single-cell rates than ammonium. For the cells assimilating these substrates, the mean single-cell assimilation rates in the Slope Site were statistically significantly higher for urea than ammonium at 1000 m, 2000 m, and 3000 m depth (Wilcoxon test, p-value <0.05), with an average of 83% higher assimilation rates for urea. The difference between urea and ammonium uptake was even more pronounced at the Open Ocean Site, with statistically significantly higher mean incorporation rates of urea at 50 m, 150 m, 2000 m, and 4000 m depth (Wilcoxon test, p-value <0.05), and an average assimilation rate an order of magnitude higher for urea than ammonium.

Assimilation of urea-derived ^13^C was also detected (Fig. S2). In the Slope site, cells assimilating urea-derived ^13^C were only detected in the epipelagic region and at 3000 m depth (up to ∼20% of cells), while in the open ocean site, cells assimilating urea-derived ^13^C were detected throughout the water column (∼25% at 500 m depth and ∼20% at 2000 m and 4000 m depth). At the Open Ocean site, we found that cell-specific ^13^C-urea assimilation was lower than that for ^15^N in the epipelagic region, but that ^13^C-urea assimilation surpassed that for ^15^N in the meso and bathypelagic regions. In general, while single-cell rates of urea-derived ^15^N assimilation showed variable trends with depth, those for urea-derived ^13^C increased with depth.

### Ammonia- and urea-based nitrification

Rates of ammonium- and urea-based nitrification were determined for all depths in the Slope and Open Ocean sites below 50 m (Figure 3). We detected the production of ^15^N-NOx in incubations with ^15^N-urea and ^15^N-ammonium at all depths in the Slope Site. Rates of nitrification were highest at 150 m (16.1 and 11.8 nmol N l^-1^ for ammonium and urea, respectively), dropped by an order of magnitude by 500 m, and then were relatively consistent with depth with a slight uptick at 3000 m. At the Open Ocean site we detected ammonium-based nitrification at 150 and 500 m, and urea-based nitrification at 150 m, but not below. Rates of urea-based nitrification were similar to that for ammonia, and statistically indistinguishable at all depths (t-test>0.1 for all depths).

**Figure 3.**
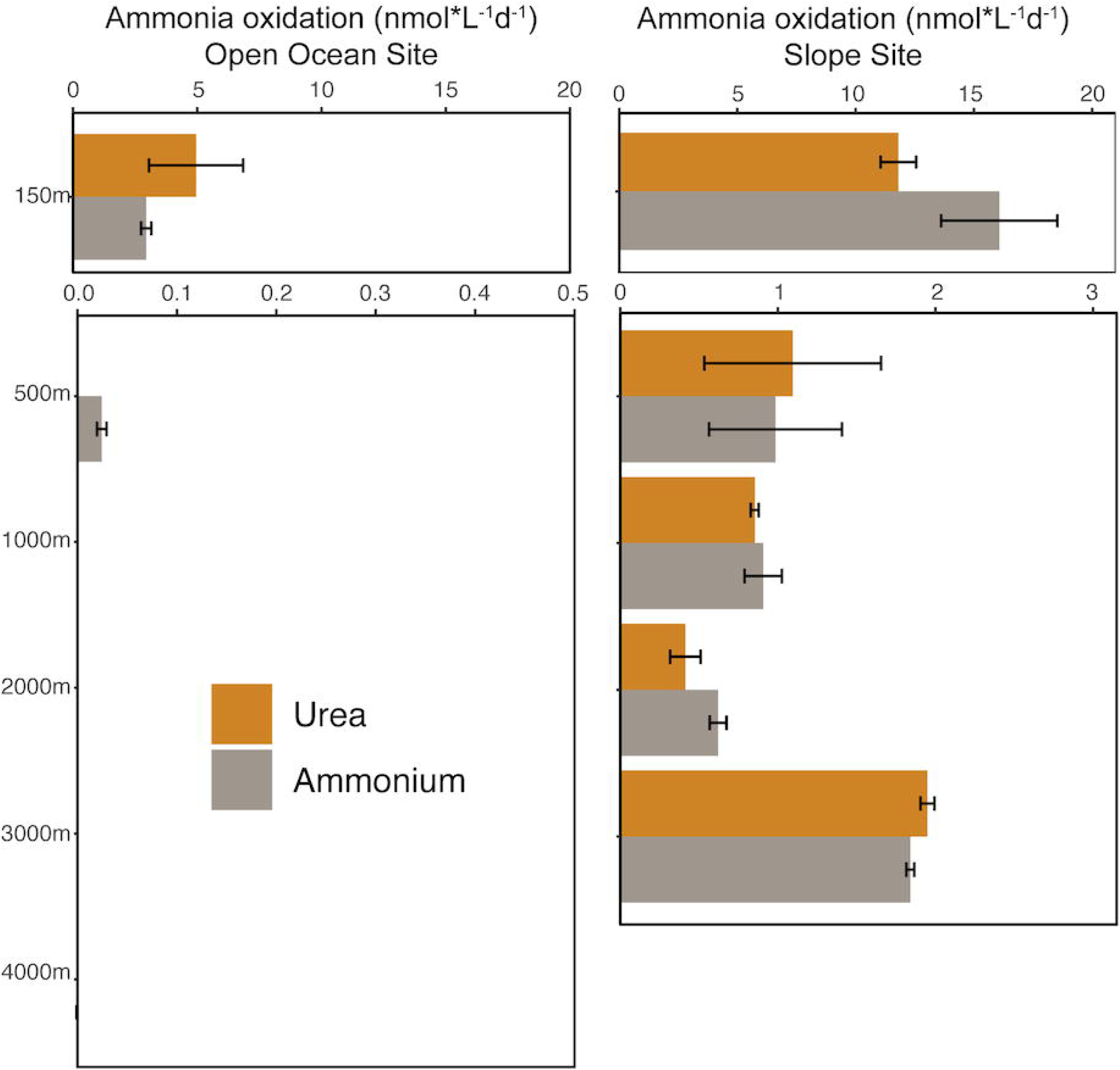
Urea- and ammonium-based ammonia oxidation rates at each depth of the Slope Site and Open Ocean Site. While at the Slope site ammonia oxidation was detected at all depths, for the Open Ocean Site no significant ammonia oxidation was detected below 150m depth (except for ammonia incubations at 500m depth). Error bars represent standard deviation of duplicate measurements.

### Relative abundance and distributions of ureC and amoA

We sequenced and assembled metagenomes (average 57.8Gb/metagenome) from all depths of the two sites where isotope incubations were conducted (Table S2), and we detected *ureC* at all depths investigated (Figure 4). The relative abundance of *ureC* genes was lowest at 50 m (0.11 *ureC*/*recA* ratio) and highest at 150 m (0.76 *ureC*/*recA* ratio) at the Slope site (Figure 4), generally mirroring the trend in proportion of cells assimilating urea at this site. The prevalence of *ureC* was more consistent with depth at the Open Ocean Site, with the maximum ratio (0.5 *ureC*/recA) found at 500 m, 1000 m, and 3000 m depth (Figure 4). We compared the relative abundance of ureC genes within the entire microbial population to that of *amoA*—the gene encoding subunit A of ammonia monooxygenase, essential for archaeal nitrification—to determine a minimum portion of ureC genes found outside of Nitrososphaeria. Trends in *ureC* and *amoA* prevalence were similar with depth, although *ureC* was consistently twice as abundant as *amoA* (0.45 *ureC*/*recA* and 0.23 *amoA*/*recA* on average; Figure 4). This indicates that even if all *amoA*-encoding organisms contained *ureC*, half of the community potential to cleave urea is found in organisms outside of that group, i.e., with different catabolisms.

**Figure 4:**
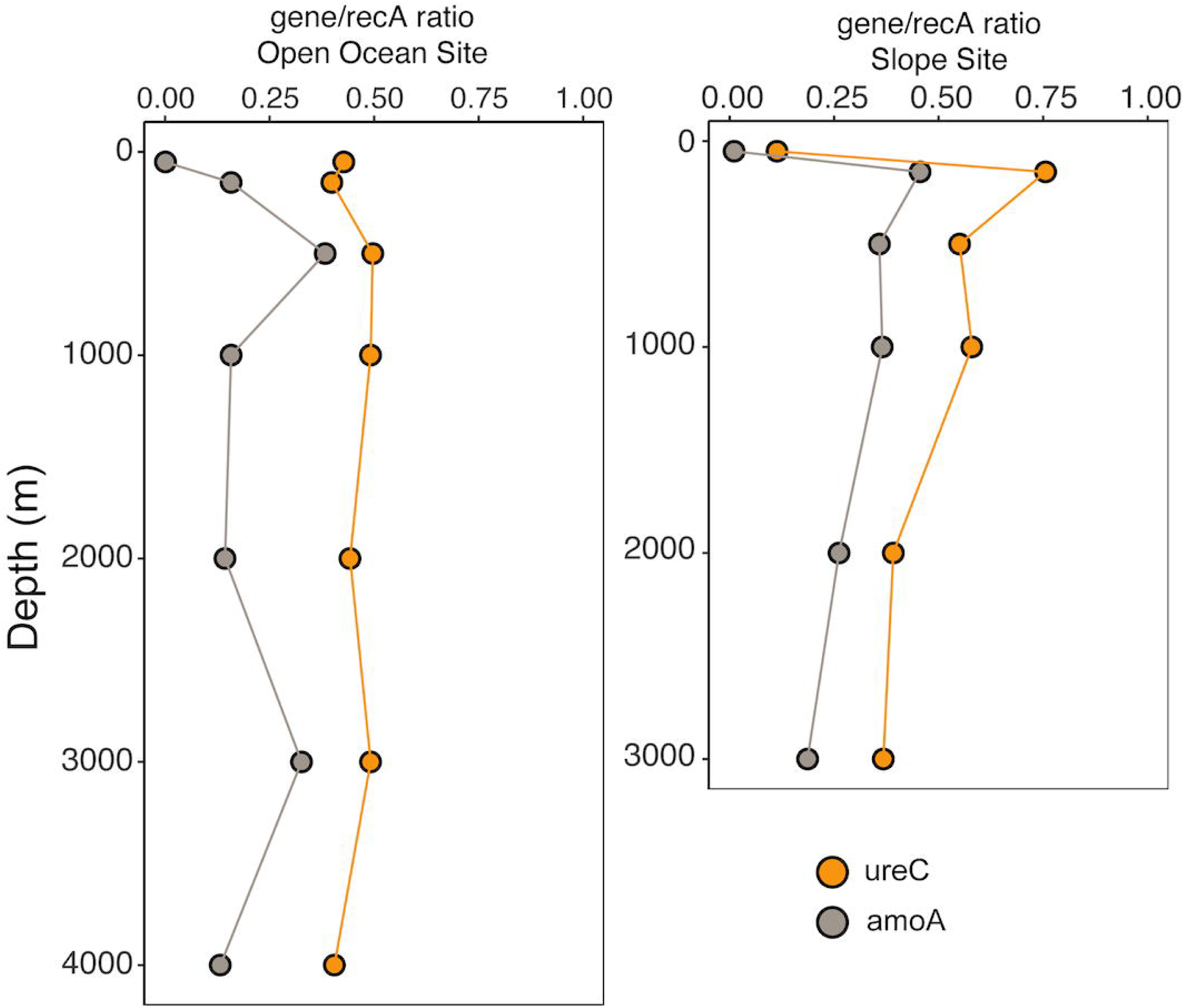
Vertical profiles showing gene abundance of *ureC* and *amoA* genes relative to a housekeeping gene (*recA*) in the metagenomes at the Open Ocean Site (left) and Slope Site (right).

### Identification and relative abundance of ureC-containing contigs

A subset of *ureC*-encoding contigs (≥3000 bp, non-eukaryotic, Table S4) were then analyzed to determine the taxonomic composition of organisms with the genetic potential to cleave urea. These *ureC*-containing contigs were associated with diverse phylogenetic groups (fourteen distinct phyla) and showed a consistent shift with depth between sites (Figure 5A, Table S3). In the 50 m sample, *ureC*-containing contigs were associated primarily with Proteobacteria at both the Open Ocean and Slope Sites (73.5% and 78.4% of the *ureC*-containing community, respectively), but were also associated with Cyanobacteria (17.7% and 10.4%), Nitrosophaerota (10.7%, only in the Slope Site), and Verrucomicrobia (7% and 0.5%). While Cyanobacteria were only detected at 50 m, members of the Nitrosophaerota comprised an increasingly large fraction of the *ureC*-containing community with depth (>50% in most samples), together with increases in members of the Planctomycetota (up to 14.0%), Verrucomicrobiota (up to 12.5%), Nitrospinota (up to 7.2%), and Myxococcota (up to 1.6%).

**Figure 5.**
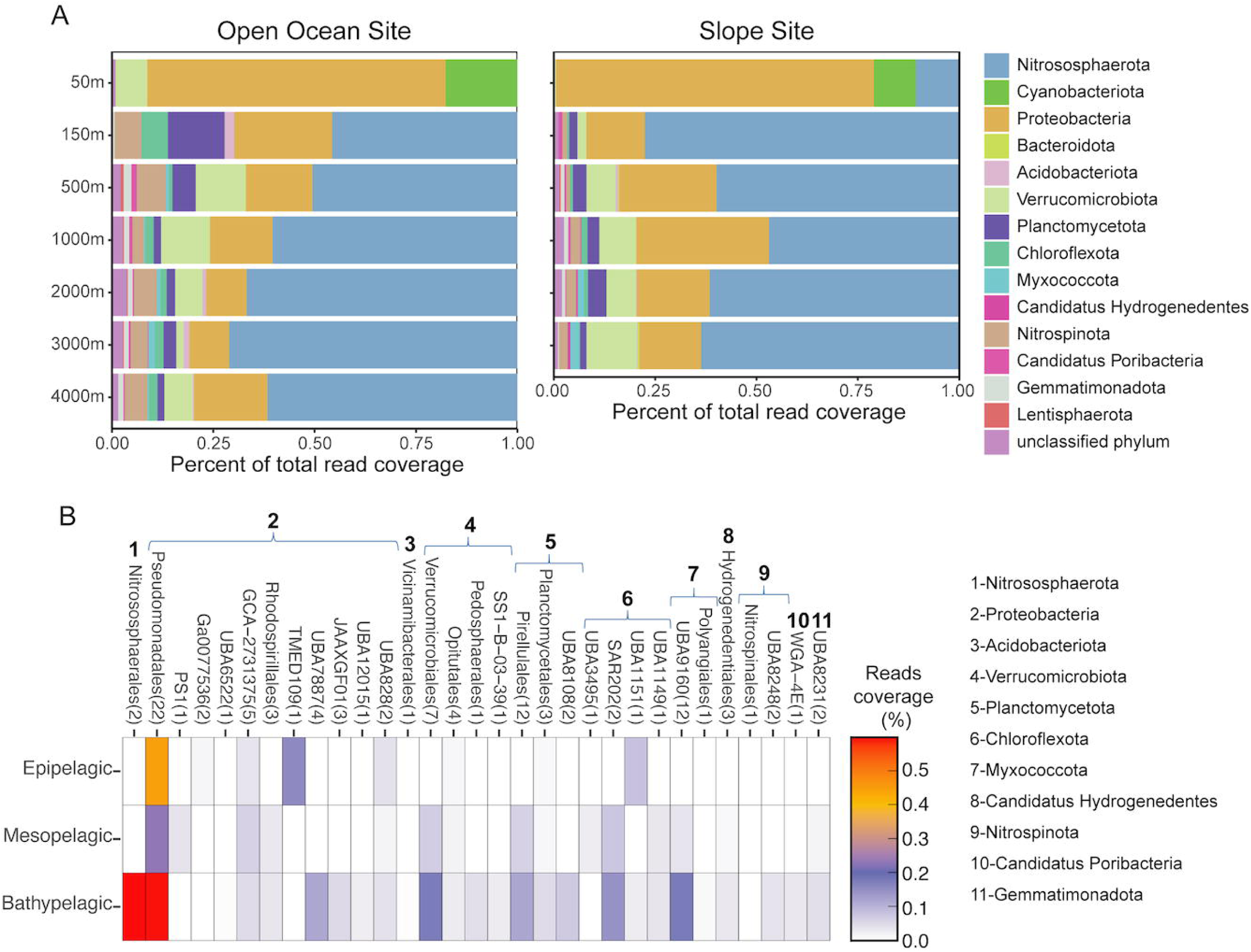
Diversity and distribution of taxa containing *ureC*. **(A)** Taxonomic analysis of the *ureC*-containing contigs classified at the phylum level for each water depth, each site separately. (B) Relative abundance of *ureC*-containing MAGs within each order (expressed as coverage of total reads) with water depth, Slope and Open Ocean sites combined. Phylum affiliation is indicated. The number of MAGs within each order is indicated in parentheses.

### Taxonomic identification and investigation of MAGs containing ureC

To complement our contig-based analyses, we generated 109 unique *ureC*-containing MAGs with >50% completeness and <5% redundancy (Table S5). Taxonomic identification of these *ureC*-containing MAGs indicated they were from twelve distinct phyla (Figure 5B), capturing most groups identified in the contig-based approach. Similarly, MAG-based abundance analyses roughly recapitulated the distribution pattern of taxonomic groups with depth, with Proteobacterial *ureC*-containing MAGs more relatively abundant at the surface and those of the Nitrosophaerota and others becoming more relatively abundant at depth. Only MAGs belonging to the Alphaproteobacterium TMED109 clade and the Chloroflexota UBA1151 were more relatively abundant in the epipelagic region than in the deepest regions. Overall, a higher relative abundance of *ureC*-containing MAG groups was found in the bathypelagic (2.56% of all mapped reads in the bathypelagic versus 0.76% in the epipelagic). MAGs within the Nitrososphaerales order (Phylum Nitrososphaerota) and the Pseudomonadales order (Phylum Proteobacteria) were the most relatively abundant ureC-containing groups in the bathypelagic region (with a coverage of 0.60% and 0.59% respectively). Other groups, including the Verrucomicrobiales and the Myxococcota UBA9160 orders, were exclusively found below the photic zone.

### UreC prevalence in global datasets and comparison to other genes

To assess the generality of our findings across the global ocean, we determined the relative abundance of *ureC* genes in epipelagic, mesopelagic, and bathypelagic depths from different ocean basins using the publicly available Tara Ocean, GEOTRACES and Malaspina databases (Figure 6, Table S6). Based on comparison with *recA*, we estimate that 10-46% of cells in the global deep sea contain *ureC* (median 36%), consistent with our findings in the North Pacific Ocean. Relative abundances of *ureC* increased with depth in the GEOTRACES data from the South Pacific[50], Tara Oceans data in the Arctic and South Atlantic Ocean[51, 52], and Malaspina data in the North Atlantic, South Atlantic, North Pacific, South Pacific, and Indian Oceans[33]. The only exception was found in sites from the northern Indian Ocean where higher *ureC* abundance was observed at epipelagic depths than mesopelagic depths (no bathypelagic metagenomes are available).

**Figure 6.**
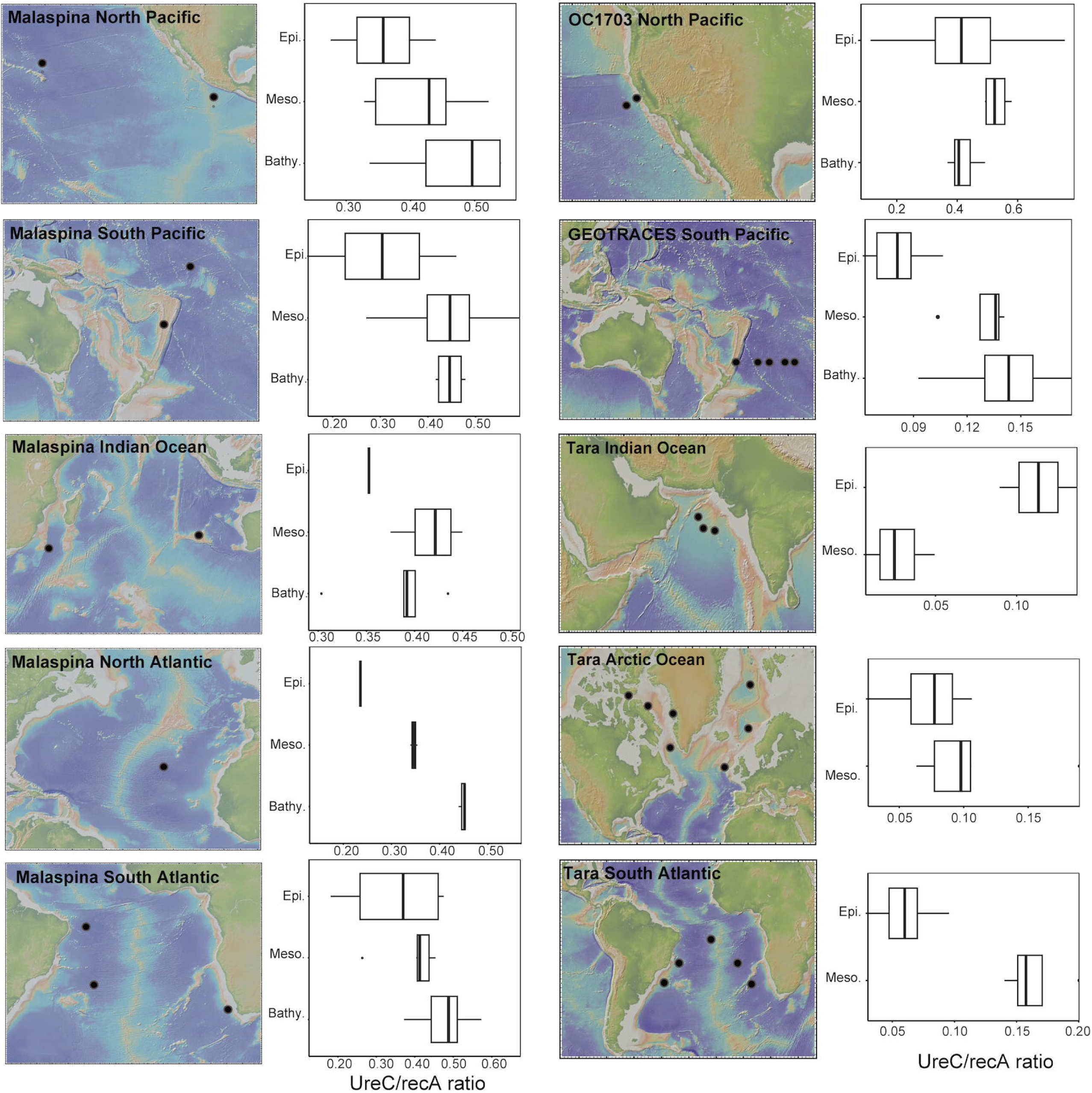
Box plots showing abundance of *ureC* genes relative to total mapped reads in the epipelagic (0-200 mbsl), mesopelagic (200-1000 mbsl), and bathypelagic (1000-4500 mbsl) regions of different ocean regions. The North Pacific Ocean is shown in the OC1703 and Malaspina datasets, Southwest Pacific Ocean in the GEOTRACES dataset, North Atlantic in the Malaspina dataset, South Atlantic and Indian Oceans in the Malaspina and Tara Oceans datasets and the Arctic Ocean in the Tara Oceans datasets. Gene abundances are displayed as the ratio between *ureC* and *recA* (both gene coverages calculated as RPKM). For the Tara Oceans, only samples for the epipelagic and mesopelagic regions were available.

We also compared the distribution of *ureC* with genes involved in the metabolism of nitrate, the most abundant source of nitrogen in the deep sea, in order to compare the potential importance of both substrates. Similar to *ureC*, genes such as *nirA* or *nasA* (key genes in assimilatory nitrate reductase), *nirB* and *nirD* (dissimilatory nitrate reduction), *nirK* and *nirS* (denitrification), as well as *nxrA* and *nxrB* (nitrite oxidoreductase) increased with depth in all analyzed datasets (OC1703, GEOTRACES and Malaspina) (Figure S3). However, *ureC* was consistently more abundant than these other genes (Figure S3). This difference is particularly notable in the Malaspina dataset, which is the dataset that best reflects the gene distribution in the world oceans, with an order of magnitude more *ureC* than *nasA*.

### Comparison of organic carbon sources to the deep sea

In order to assess the significance of deep-sea nitrification to the marine carbon cycle we estimate and compare two sources of organic carbon to the deep sea: the particulate organic carbon (POC) consisting of photosynthetic detritus sinking from the euphotic zone (‘gravitational POC’) and the inorganic carbon fixed via nitrification-based chemoautotrophy in the deep sea. We estimated POC export fluxes to be 26 and 128 mg C/m^2^/day at the base of the euphotic zone (100 m) at the open ocean and coastal sites, respectively. In parallel, assuming a DIC fixation yield of 0.09 mol C fixed/mol N oxidized[53], we converted the rates of urea- and ammonium-based nitrification measured here into carbon fixation rates and integrated them over the dark water column (100-3000 m or 100-4000 m, depending on the site). We found rates of 1.4 and 8.8 mg C/m^2^/day of autotrophic carbon fixation for the open ocean and slope sites respectively, corresponding to 5 and 7% of the gravitational POC fluxes entering the dark ocean.

## Discussion

Urea is increasingly recognized as a source of nitrogen for cell growth [30] [20, 54] and nitrification[17, 18] in the sunlit ocean. In the euphotic zone, nitrogen from urea is assimilated by phylogenetically diverse taxa, including Cyanobacteria, Proteobacteria, and Thermoproteota (e.g., Nitrosophaeria) [14, 18, 20] [19, 28, 55], at rates exceeding those for nitrate, leucine, glutamate[14], and even ammonium[56], and it is also oxidized by nitrifying Nitrosophaeria to support chemoautotrophy[57, 58]. Our observations in the aphotic zone of the northwest Pacific Ocean indicate that the significance of this molecule not only extends to the aphotic zone— where an equally broad yet predominantly different set of organisms cleave it—but indicate that its role may be even more central to ecosystem functioning there than in surface waters. Our data show that its availability—assessed as a function of both its concentration and the prevalence of the genetic capacity to access it—increases with depth, as does the microbial preference for it as a nitrogen source over ammonia. Additionally, while rates of urea-based nitrification are lower at depth than at the surface, they are comparable to that of ammonia in both realms, and likely play an outsized role in microbial community dynamics at depth by supporting the production of organic matter in a more energy- and carbon-limited system than at the surface.

Urea-derived nitrogen is widely and extensively assimilated by microrganisms in the aphotic zone. We find that on average 25% of cells in the meso- and bathypelagic assimilate urea-derived nitrogen. Assuming that ammonium uptake reflects cellular activity[7] and then comparing the subset of cells assimilating urea-N to that assimilating ammonium-N, we find that 60% of active cells in the meso- and bathypelagic assimilate urea-derived nitrogen. Although cross-feeding of ^15^N-labelled substrates can cause these proportions to be greater than the number of cells directly consuming urea, the paired metagenomic data is roughly consistent with these values in the deep-sea; we estimate that an average of 39% of cells in the meso- and bathypelagic at these sites contain a *ureC* gene, putatively allowing lineages from at least fourteen distinct phyla to cleave urea directly (as described in more detail below). Additionally, cells assimilating urea-derived N did so at higher rates than cells assimilating ammonium in almost all samples investigated. This demonstrates the significance of this nitrogen source to the cells and also decreases the likelihood that cells assimilated the urea-derived N as recycled ammonium. Regardless of what proportion of the assimilation was directly from urea versus recycled substrates, the widespread and high rates of consumption of urea-derived N indicates that the large reservoir of urea-nitrogen in the deep sea—on average an order or magnitude more abundant than ammonium—is available to most cells.

Nitrate remains the largest pool of fixed nitrogen in the deep sea, averaging over two orders of magnitude more abundant than urea at our site. Our observations of urea assimilation occurred in the presence of these high concentrations of nitrate, suggesting a preference for urea over nitrate. Indeed, we found that *ureC* genes were more abundant than those related to assimilatory nitrate reduction (such as *nasA*) within our study sites, as well as a broad distribution of publically available deep sea datasets (Fig. S3). Previous work has also reported relatively low detection of *nasA* in the Malaspina global deep-sea metagenomic dataset [6, 7]. Preference for urea is likely related to the higher energy requirements of the assimilatory reduction of nitrate [59][60], a difference that might be particularly relevant in the energy-poor aphotic zone. While gene abundances are useful indicators of potential activity, and how well a given ability is distributed across a community, direct comparisons of the uptake of nitrate and urea in the deepest regions of the oceans would be beneficial to directly compare the proportions of cells capable of assimilating each, and with what preference. Notably, the meso- and bathypelagic regions accounted for nearly half of the total pelagic urea assimilation in the Slope Site—more than it contributed to either ammonium or amino acid assimilation [7]—indicating that urea is a more important nitrogen source in the deep sea relative to the surface than for either ammonium or amino acids.

Urea also represents a major potential substrate for nitrification by members of the Nitrosophaeria phylum [17–19, 25]. Remarkably, urea-based nitrification can also happen in the presence of ammonium[18], suggesting that urea is not only an alternative for ammonium when it is scarce, but can also represent a primary substrate for nitrification. Furthermore, a recent study shows that some ammonia-oxidizing bacteria repress the use of extracellular ammonium in the presence of ammonium derived from urea hydrolysis in the cytoplasm[56]. While previous studies have highlighted the significance of urea-driven nitrification in the epipelagic[17, 18, 61] and upper mesopelagic region (up to 300 m depth[25]), as well as demonstrated the potential of deep marine microbes to oxidize ammonia through genomic analysis[19], urea-driven nitrification has not been directly measured in the lower mesopelagic and bathypelagic regions. The detection of urea-based nitrification at all depths of our Slope site suggests that deep-sea nitrifiers can indeed use urea as a substrate (Figure 2). Oxidation of ammonia after urea hydrolosis by other community members is also possible, but regardless, this confirms urea-derived nitrogen is readily available to microbes for nitrification. Direct oxidation of urea by deep-sea nitrifiers is additionally supported by the metagenomic analysis, which showed not only that *ureC* genes were found within Nitrosophaeria MAGs, but that over half of read coverage of *ureC*-containing contigs in the meso- and bathypelagic was affiliated with Nitrososphaeria (Figure 5A). The rates of urea-based nitrification were statistically indistinguishable from those for ammonium at all depths, consistent with the previous work at 300 m depth[25], indicating a potentially significant role for urea in deep-sea nitrification. Rates of both ammonia- and urea-based nitrification decreased with depth and distance from shore (Figure 2), consistent with the trends we observed in overall anabolic activity previously at this site [7].

Using metagenomics, we determined both the distribution of *ureC* in microbial communities at our study site and in globally-sourced datasets, and also phylogenetically identified the taxa containing *ureC*. We detected *ureC* genes throughout the water column, and found that their prevelance—the proportion of microbial cells possessing it in a given sample— reached a maximum in the aphotic zone in both our study site and the other global datasets we analyzed. Overall, we see that about a third of the cells in the dark ocean (average 39% in our dataset, and average of 30% in the global datasets) contain *ureC*. Both the contig- and MAG-based analyses identified diverse taxa containing *ureC* genes at our site, with fourteen distinct phyla identified by the former and twelve by the latter. While the MAG-based phylogenetic identification of *ureC*-containing genomes is likely more robust than the contig-based identifications, due to the greater sequence length available for consideration, the contig-based approach provides a more comprehensive overveiew of the community, including taxa that may systematically evade genomic binning. Notably, twelve of the fourteen phyla identified in the contig-based analysis were also identified with the MAG-based analysis. The groups identified as containing *ureC* in the 50 m samples are generally consistent with previous work in the euphotic zone, especially in the identification of Gammaproteobacteria and Prochlorococcus[14].

The deep-sea analysis revealed that some taxonomic groups with members known to use urea at the surface also have members with the genetic capacity to do so at depth, including Nitrososphaerales, Verrucomicrobiota, and Myxococcota, as well as members of several groups not before reported to utilize urea, including SAR202 and Alphaproteobacteria TMED109. We interpret the presence of *ureC* genes in taxa not known to oxidize ammonia, and in MAGs without an *amoA* gene, as evidence of potential urea use for nitrogen acquisition. When found together with *amoA* (i.e., within the Nitrosphaeria), it may be used for both nitrogen acquisition for biomass and for nitrification. While our metagenomic analysis is consistent with a large role for urea in nitrifying organisms in the deep sea – evidenced by the large fraction of *ureC* genes within the Nitrosphaeria – our work also highlights the wide diversity of organisms capable of cleaving it. And, as not all nitrifiers contain urease (e.g., *Nitrosopelagicus brevis* CN25, Santoro et al., 2015, *Nitrosopumilus maritimus*[23]), there may be an important relationship between heterotrophic urea degraders and the chemoautotrophic nitrifiers, with ammonia shared in one direction and organic carbon in the other.

The implications of deep-sea nitrification on the marine carbon cycle are considerable. It is often assumed that the main—and essentially only—source of organic carbon to the dark ocean is gravitational POC[62]. However, the persistent imbalance between known supply and demand of organic matter in the deep sea highlight the inadequacies of current knowledge[63]. Our observations indicate that urea- and ammonium-based nitrification could together provide 5 and 7% of the estimated gravitational POC entering the top of the mesopelagic at the Open Ocean and Slope Sites, respectively. These are significant values considering gravitational POC fluxes decrease logarithmically with depth in the water column[64, 65]. Additionally, while organic carbon generated at depth is generally labile, gravitational POC is increasingly recalcitrant and biologically inaccessible with depth, highlighting the potential importance of even small amounts of endogenously produced organic carbon[66]. To fully assess the role of chemoautotrophy in helping balancing the carbon budget more studies are required, including direct measurements of carbon fixation ideally at *in situ* pressures and concurrent measurement of sinking POC flux. However, our estimates provide new evidence that deep-sea chemoautotrophy, which is often overlooked in models of the biological carbon pump, is not negligible, and could be of paramount importance.

In summary, our study reveals a large reservoir of urea-N in the deep sea (Figure 1), widespread genetic potential for urea utilization in the meso- and bathypelagic (Figures 4 and 5), and direct evidence for both extensive assimilation of urea-derived nitrogen (Figure 2) and the persistence of both urea- and ammonia-based nitrification throughout the epi-, meso-, and bathypelagic (Figure 3). While additional direct measurements are necessary to confirm our results globally, we contend that urea use is likely widespread throughout the global deep sea on the basis of the generally physicochemically representative nature of our study site and the high proportions of *ureC*-encoding microorganisms throughout the global metagenomic datasets analyzed here. These results address long-standing hypotheses about the potential for urea to fuel nitrification in deep waters, and indicate the potential for chemoauototrophy at depth to significantly impact the marine carbon budget.

## Data Availability

Read data and metagenome-assembled genomes (MAGs) analyzed in this study are available through NCBI at PRJNA1054206.

## Acknowledgements

We thank Julian Fortney, and Nicolette Meyer, as well as the captain and crew of R/V Oceanus cruise OC1703A, for assisting with field sampling. This work was primarily supported through a Simons Foundation Early Career Investigator Award to AED (507798 ) and a National Science Foundation CAREER Award to AED (2143035). Cruise OC1703A was supported by NSF Award 1634297 to AED. The nanoSIMS analyses were performed at the Stanford Nano Shared Facilities (SNSF), which is partially supported by the National Science Foundation under award ECCS-2026822. We thank Christie Jilly-Rehak and Chuck Hitzman for assistance with the nanoSIMS analyses. We thank Pascale Anabelle Baya-Ardyna for assistance with the nitrification measurements. NAG was supported by the ‘Severo Ochoa Centre of Excellence’ accreditation (CEX2019-000928-S) funded by AEI 10.13039/501100011033, and the Beatriu de Pinós program (2020-BP-00179) during the writing of this manuscript. ALJ was supported by the Stanford Science Fellows program and the National Science Foundation Postdoctoral Research Fellowship in Ocean Sciences. RSRS was supported by the Stanford Graduate Fellowship Program.

## Conflicts of Interest

None to declare.

## Notes

### Competing Interest Statement

The authors have declared no competing interest.

