## Supplementary Text/Figures for "Urea assimilation and oxidation supports the activity of a phylogenetically diverse microbial community in the dark ocean"

**Supplementary Information**

Supplementary Text: Supplementary Materials and Methods

Supplementary Tables 1-7 captions. The tables are provided as a separate single xlsx file.

Supplementary Figures 1-3

**Supplementary Materials and Methods**

*NanoSIMS analysis conditions*

Portions of the filter samples were mounted onto conductive carbon tabs and coated with gold in a sputter coater. Areas of filters were randomly selected for the analysis using a charge-coupled device (CCD) camera. A NanoSIMS 50L (CAMECA, Gennevilliers, France) housed in the Stanford Nano Facility was used with a primary beam of $Cs^+$ ions operating in continuous mode. The beam was set up at 4 pA and samples were analyzed using a 256 x 256 pixel raster on 30 x 30 μm areas with~1 ms/pixel dwell time. 30-second pre-sputtering with a high energy beam was used to partially remove the gold coat on top of the cells. The masses collected with electron multipliers were $^{12}C^-$, $^{13}C^-$, $^{12}C^{13}C^-$, $^{12}C^-_2$, $^{12}C^{14}N^-$, $^{12}C^{15}N^-$ and $^{32}S^-$. Forty-five serial secondary ion images (i.e., layers) were collected at each analysis area. The mass resolving power (MRP) was ~7000 (after x1.5 correction[1]) with an exit slit width of 70 μm, an entrance slit width of 20 μm, an aperture slit of 200 μm, and the energy slit filtering about 30% of secondary ions from the high-energy tail of their energy distribution.

*Metagenomic pre-processing, assembly, and gene-based analysis*

Read trimming via bbduk employed the following parameters: *ktrim=r k=23 mink=11 hdist=1 tbo qtrim=r trimq=25 minlen=20*. Assemblies of each individual sequencing dataset by MEGAHIT employed the *meta-sensitive* preset. Abundance of *ureC* within assemblies was analyzed as described in the main text. To analyze the abundance of the rest of the genes, including *recA*, *amoA*, and nitrate-related genes (*nasA*, *nirA*, *nirB*, *nirD*, *nirK*, *nrxA*, and *nrxB*), gene prediction was carried out by Prodigal[2] and gene annotation was carried out using KofamScan against the KEGG database. As before, trimmed metagenomc reads were mapped against identified genes. RPKM coverage values for each gene were calculated using the number of mapped reads, total mapped reads for the sample, and average gene length. The abundance of *urec* and other genes was displayed as the ratio of the gene RPKM and the *recA* RPKM.

*Taxonomic composition of ureC-encoding contigs*

To examine the full phylogenetic diversity of organisms encoding urease, we extracted nucleotide sequences from all OC1703 contigs predicted to encode *ureC*. Protein sequences for all open reading frames (ORFs) on these contigs were predicted using Prodigal and aligned against UniRef100[3] using DIAMOND (*blastp, -b8, -c1)*[4]. Alignments were filtered to those with an e-value ≤1e-20 and a query coverage ≥70%, and the top-scoring alignment per ORF was retained. UniRef/UniParc identifiers were extracted and mapped to NCBI taxonomic strings using the batch ID mapping tool on uniprot.org and the NCBI taxonomy database (https://www.ncbi.nlm.nih.gov/taxonomy). For each *ureC*-encoding contig, a preliminary consensus taxonomy was assigned by taking the most common phylum-level designation predicted by protein alignments, as has been done previously[5]. Taxonomic assignments for all contigs ≥3000 bp in length were manually curated, with an emphasis on those contigs < 5kb or with a weak taxonomic signal. Manual curation was aided by grouping contigs into sequence-based clusters using dRep *(-pa 0.80 -sa 0.95 --clusterAlg single*)[6] and by comparing predicted contig taxonomy to bin taxonomy, where contigs were able to be binned (see below section). Sequence fragments predicted to derive from eukaryotic genomes were discarded.

*Binning and analysis of ureC-encoding genomes*

To complement our contig-based analyses of *ureC*-containing organisms, we subjected our metagenomic assemblies from the Open Ocean site (OC6) and the Slope Site (OC3) to binning. Assemblies from these sites (n=13) were used to create index files against which trimmed metagenomic reads from all sites were aligned using bowtie2[7]. Contig coverage tables were created using the jgi_summarize_bam_contig_depths script from MetaBAT2[8] and binned using a minimum contig size of 1500 bp. Resulting bins were assessed for quality using CheckM[9] and taxonomic classification using GTDB-Tk (ver. 2.3.2)[10]. Those MAGs encoding the urease genes were identified by cross-referencing member contigs with the list of *ureC* genes identified in the contig-based analyses (see section above). Secondary refinement of MAGs encoding urease was performed using Anvi'o[11], in which contigs with anomalous coverage and GC profiles were removed. After refinement, CheckM quality metrics were recomputed. Finally, the relative abundance of the refined MAGs from different taxonomic lineages was computed by coverm (version 0.6.1).

**Supplementary tables captions:**

**Supplementary table 1:** Summary of sample locations and environmental data, including temperature, salinity, fluorescence, dissolved oxygen, and concentration of different nitrogen species.

**Supplementary table 2:** Characteristics of metagenomic samples analyzed in this study. perc_reads_mapping describes the percent of trimmed metagenomic reads mapping to the assembly with loose mapping parameters.

**Supplementary table 3:** Features of ureC-encoding contigs of the OC1703 dataset used to analyze the taxonomic composition of urea utilizers across depth.

**Supplementary table 4:** Features of the OC1703 dataset contigs containing ureC, amoA, or recA used to estimate gene coverage.

**Supplementary table 5:** Characteristics of metagenome-assembled genomes (MAGs) analyzed in this study.

**Supplementary table 6:** Features the ureC-containing contigs for the Tara Oceans, GEOTRACES, and Malaspina datasets.

**Supplementary table 7:** ureC Gold Standard sequences used as BLAST query sequences against Tara Oceans OM-RGC_v2_assemblies and for Transitivity Clustering.

**Supplementary figures:**

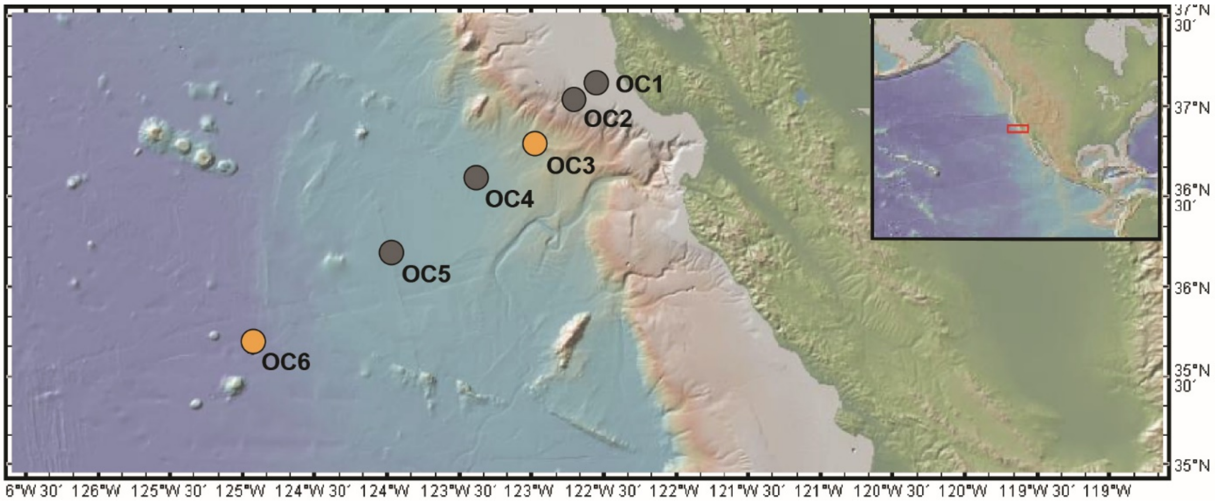


**Supplementary Figure 1. (A)** Map of the northeast Pacific Ocean showing the location of the sampling sites. The sites colored orange indicate the locations where samples for nanoSIMS and metagenome analysis were taken. Map was generated with GeoMapApp Version 3.6.15.

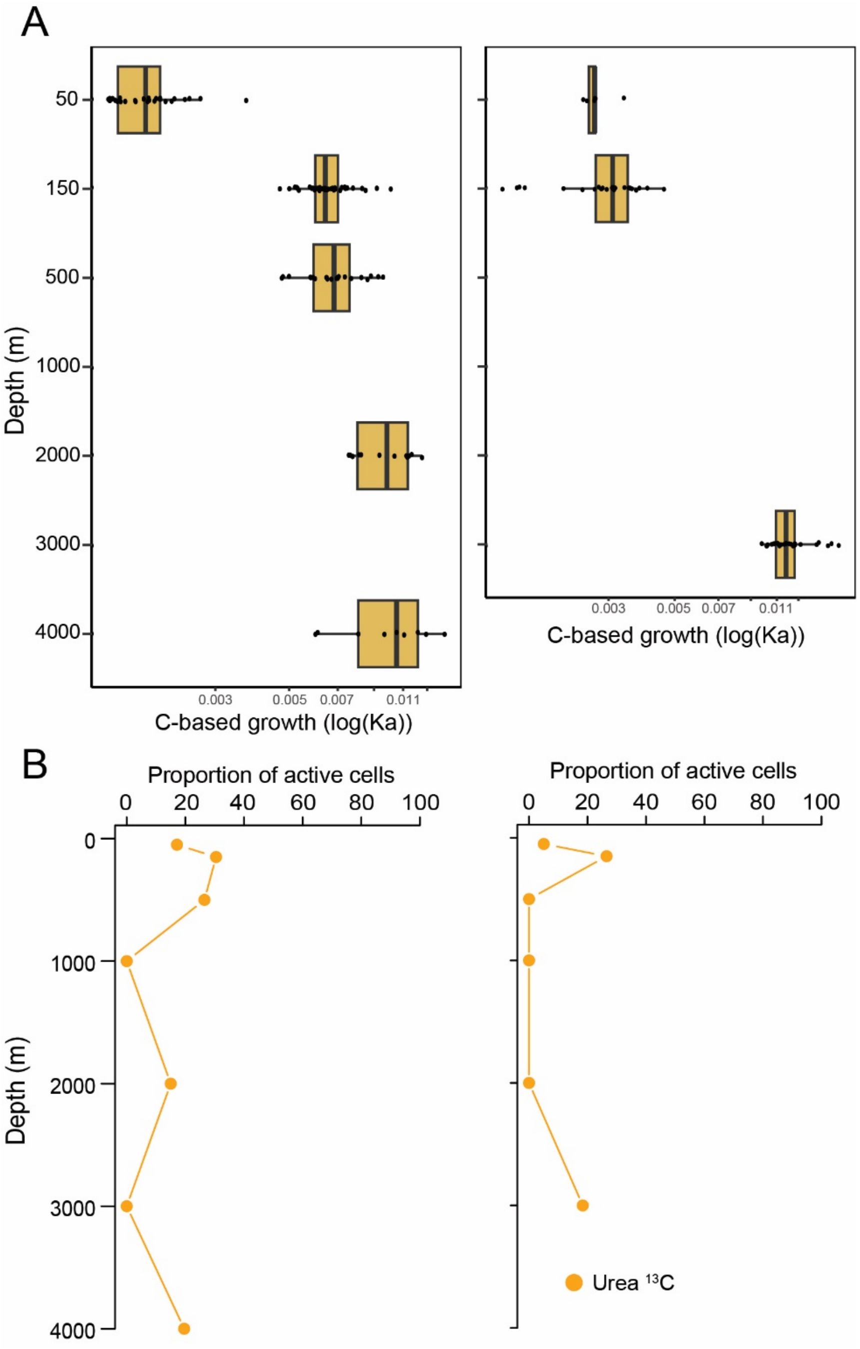


**Supplementary figure 2.** (A) Boxplots representing the assimilation rates (Ka) in logarithmic scale for $^{13}C$ urea in the Open Ocean Site (left) and Slope Site (right). (B) Proportion of active cells assimilating $^{13}C$-urea of all cells analyzed in the Open Ocean Site (left) and Slope Site (right).

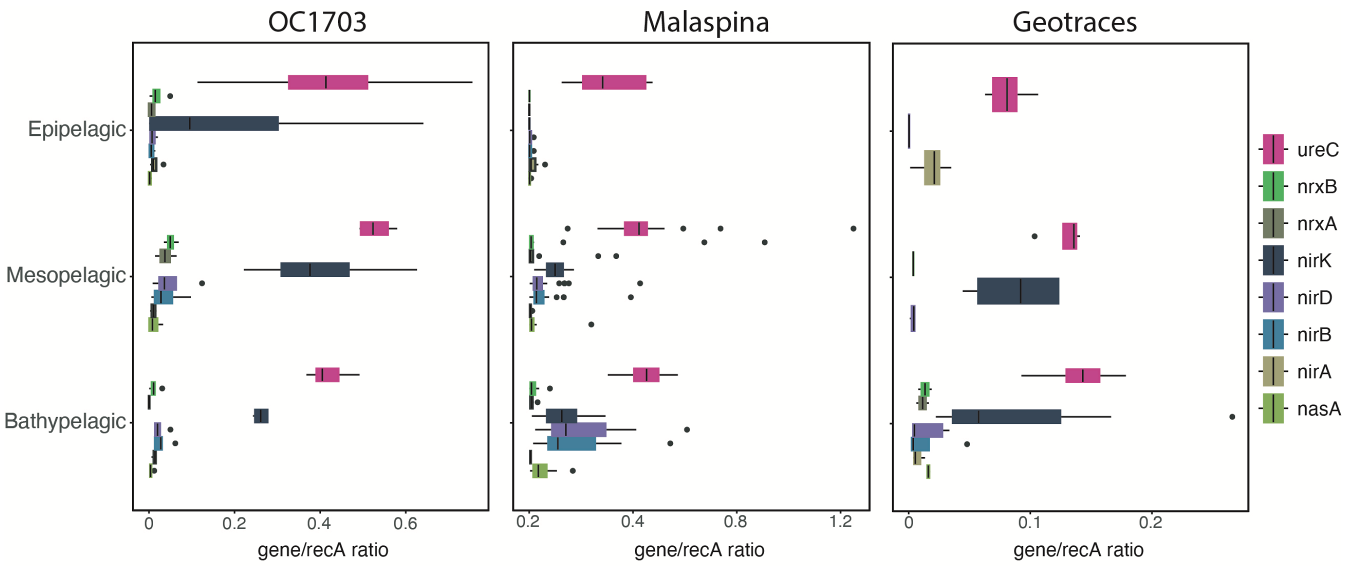


**Supplementary figure 3.** Boxplot showing the relative abundance of ureC gene, as well as main genes involved in assimilatory nitrate reduction, dissimilatory nitrate reduction, and denitrification. Boxplot was built using the entire Malaspina metagenome dataset for the global epipelagic, mesopelagic and bathypelagic regions and they were estimated using the single copy gene normalized abundance.
